# Does relaxing the infinite sites assumption give better tumor phylogenies? An ILP-based comparative approach

**DOI:** 10.1101/227801

**Authors:** Paola Bonizzoni, Simone Ciccolella, Gianluca Della Vedova, Mauricio Soto

## Abstract

Most of the evolutionary history reconstruction approaches are based on the infinite site assumption, which is underlying the Perfect Phylogeny model and whose main consequence is that acquired mutation can never lost. This results in the clonal model used to explain cancer evolution. Some recent results gives a strong evidence that recurrent and back mutations are present in the evolutionary history of tumors [5,21], thus showing that more general models then the Perfect Phylogeny are required. We propose a new approach that incorporates the possibility of losing a previously acquired mutation, extending the Persistent Phylogeny model [1].

We exploit our model to provide an ILP formulation of the problem of reconstructing trees on mixed populations, where the input data consists of the fraction of cells in a set of samples that have a certain mutation. This is a fundamental problem in cancer genomics, where the goal is to study the evolutionary history of a tumor. An experimental analysis shows the usefulness of allowing mutation losses, by studying some real and simulated datasets where our ILP approach provides a better interpretation than the one obtained under perfect phylogeny assumption. Finally, we show how to incorporate multiple back mutations and recurrent mutations in our model.

## 1 Introduction

Character-based phylogeny reconstruction is one of the fundamental problems in Bioinformatics, with a large literature [12,15,28,30] focusing on a simple assumption: the input data consists of a set of species (or individuals) for which we know the set of characters that it possesses. In this case, the goal is to compute a phylogeny that explains the set of input species and characters, where each edge of the phylogeny allows characters gains and losses. Character-based phylogenies play a crucial role in modeling the evolution in cancer genomics. Cancer is an uncontrolled evolutionary process of somatic mutations of tumor cells from a single founder cell [13] creating a diverse set of subpopulations [8,22,31], each originated from a single *clone*: each clone (and each subpopulation) has a distinctive set of mutations. From this point of view, a tumor progression is a phylogeny where clones and mutations have the same role as species and mutations in the classical phylogeny reconstruction setting as characters.

To fall within the classical framework we would need to obtain data directly from a cell. Unfortunately, single cell sequencing is not cheap [24] and is prone to errors, therefore we have to study *samples* comprising lots of cells belonging to an unknown set of subpopulations. This adds a new complication, since for each sample we know the (approximate) fraction of cells that have a given somatic mutation. More precisely, each read extracted from the sample is mapped against the reference genome, therefore we obtain the mutations of each read. Errors in read, repeated regions of the genome, and the fact that the coverage of the reads is not uniform throughout the genome or the cells of the sample, means that the fraction of reads that have a mutation is only an approximation of the fraction of cells of the sample that have that mutation. In other words, the observed frequencies are an estimate of the true frequencies of the cells that have a mutation.

The above reasoning leads to a computational problem called *variant allele frequency factorization problem* (VAFFP) [9,10,18], where the input is the observed frequencies of the mutations in each sample and the desired output is a phylogeny representing the tumoral evolution, as well as the composition of each sample in terms of the subpopulations or clones. The literature has mainly focused on the infinite site assumption [9], that is also known as perfect phylogeny [15], where samples contain mixtures of two-state characters, i.e. (1) each character/locus is either mutated or not, and (2) each mutation can be gained only once and never lost in the entire history of the tumor.

A possible generalization (that we do not explore in this paper) is the multi-state perfect phylogeny that has been recently proposed in order to take into account the effect of copy number aberrations on alleles [10]. In this new model — known as the infinite allele assumption — the characters can assume different states (*i.e.*, the number of copies of a site) but, as in the binary case, a change to a given state can occur only once. This restriction allows to obtain efficient algorithms, but most recent studies refutes it [21] and state that more complex models are needed to describe the tumor evolution. More precisely, deletion of entire genome regions are quite common in tumors, therefore a mutation is acquired only once, but can be then lost, even more than once. In this paper we describe an ILP-based approach that overcomes this limitation and allows to reconstruct phylogenies capturing a likely evolutionary history of the tumor studied.

We will focus on three main character-based models that generalize the Perfect Phylogeny: the Persistent Phylogeny [1] (where each character can be gained once and lost at most once), the Camin-Sokal [6] (where each character can be gained several times, but never lost), and the Dollo [11] (where each character can be gained at most once, but lost several times). We denote by Camin-Sokal(*k*) the restriction of the Camin-Sokal model where each character can be gained at most *k* times in the entire tree. Moreover, we denote by Dollo(*k*) the restriction of the Dollo model where each character can be lost at most *k* times in the entire tree. Clearly, the Persistent Phylogeny [1] corresponds to the Dollo(1) model which has been recently investigated in several works aiming to develop efficient solutions for the model [3,4,16] since its use is motivated also in other contexts [2,26]. In particular, in [1] it is proved that the Persistent Phylogeny Problem over a binary matrix *M* can be formulated as finding a special completion of an extended matrix *M_e_* that is a Perfect Phylogeny. Based on this characterization, an ILP formulation for the Persistent Phylogeny has been developed in [16].

In [9] the approach used to solve the VAFFP problem is a combination of an integer linear programming (ILP) formulation and a clever approach to compute the set of relevant phylogenies, based on the notion of ancestry graph. Since the last component is tightly coupled with the fact that perfect phylogenies have as many species as characters, it is not immediate to extend the approach of [9] to more general models. Another approach to solve the problem is based on quadratic integer programming [23], but this technique is unlikely to scale to larger datasets: for this reason the authors also provide a heuristic.

We combine some of the main ideas of the ILP formulations of [9,16] with the characterization in [1], to design a novel approach to the VAFFP problem that is entirely based on ILP and allows to take into account the three evolutionary models presented above. We have analyzed experimentally our ILP approach on both simulated and real data to test if our approach is applicable in practice as well as whether allowing the models to violate the infinite site assumption leads to better predictions. Indeed, our experiments show that the Persistent phylogeny that we compute usually provides a better interpretation of the input data than the Perfect Phylogeny, by computing a phylogeny with smaller overall error. while requiring a number of clones that is smaller than the number of mutations. Finally, the inferred tree from real data on a Leukemia tumor CLL077 reveals the losses of a mutation, though being the tree mostly consistent with the one reconstructed by other known methods [20].

## 2 Preliminaries

### 2.1 The Variant Allele Frequency Factorization Problem

The input of our main problem is a *p* × *m frequency* matrix *F* which contains the frequencies of the mutation in a set of samples. Namely, each entry *F*[*t*, *j*] indicates the proportion of cells in sample *t* having the mutation *j*. A *p* × *n usage* matrix *U*, contains the mixture of cells in each sample. More precisely, each entry *U*[*t*, *i*] is the proportion of the cells in the sample *t* belonging to the subpopulation *i*. Finally, the *n* × *m* (clonal) matrix *M* contains which subpopulation has a given mutation. An evolution model *𝓜* consists of a set of constraints that a phylogeny *T* realizing the clonal matrix *M* must obey. For example, when the evolution model is the persistent phylogeny, then the phylogeny *T* cannot have two edges corresponding to two gains or two losses of the same character. The 𝓟-VAFF problem can be formally defined as follows.

#### Definition 1

Given a *p* × *m* frequency matrix *F*, a number of clones *n*, and an evolution model 𝓟, the 𝓟-VAFFP (short for 𝓟-Variant Allele Frequency Factorization Problem) asks for an *p* × *n* usage matrix *U* and an *n* × *m* clonal matrix *M* such that (1) 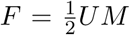, and (2) *M* admits a phylogeny under the model 𝓟.

The 1/2 factor in the definition is a technical consequence of the fact that the healthy (wild type) cell subpopulation exists, but is not one of the clones of *M*, that human beings are diploid, that is they have two copies of each chromosome, and that mutations are acquired rarely, so only one of the two copies is affected.

The 𝓟-VAFFP problem, when 𝓟 is a perfect phylogeny was first introduced in [9]. This formulation is heavily based in the infinite sites assumption which implies that no two mutations can happen at the same site. Consequently, the evolutionary history consists of a mutation gains that can be represented as an ancestry relation between mutations. Then the VAFFP problem can be reduced to a restricted version of the spanning tree problem. Furthermore, we stress the fact that in this setting the number of clones must equal the number of mutations of the frequency matrix. This fact does necessarily hold more general evolution models are used, since the infinite site assumption can be (and usually is) violated and there is no 1-to-1 mapping between edges of the tree and mutations.

**Figure 1:**
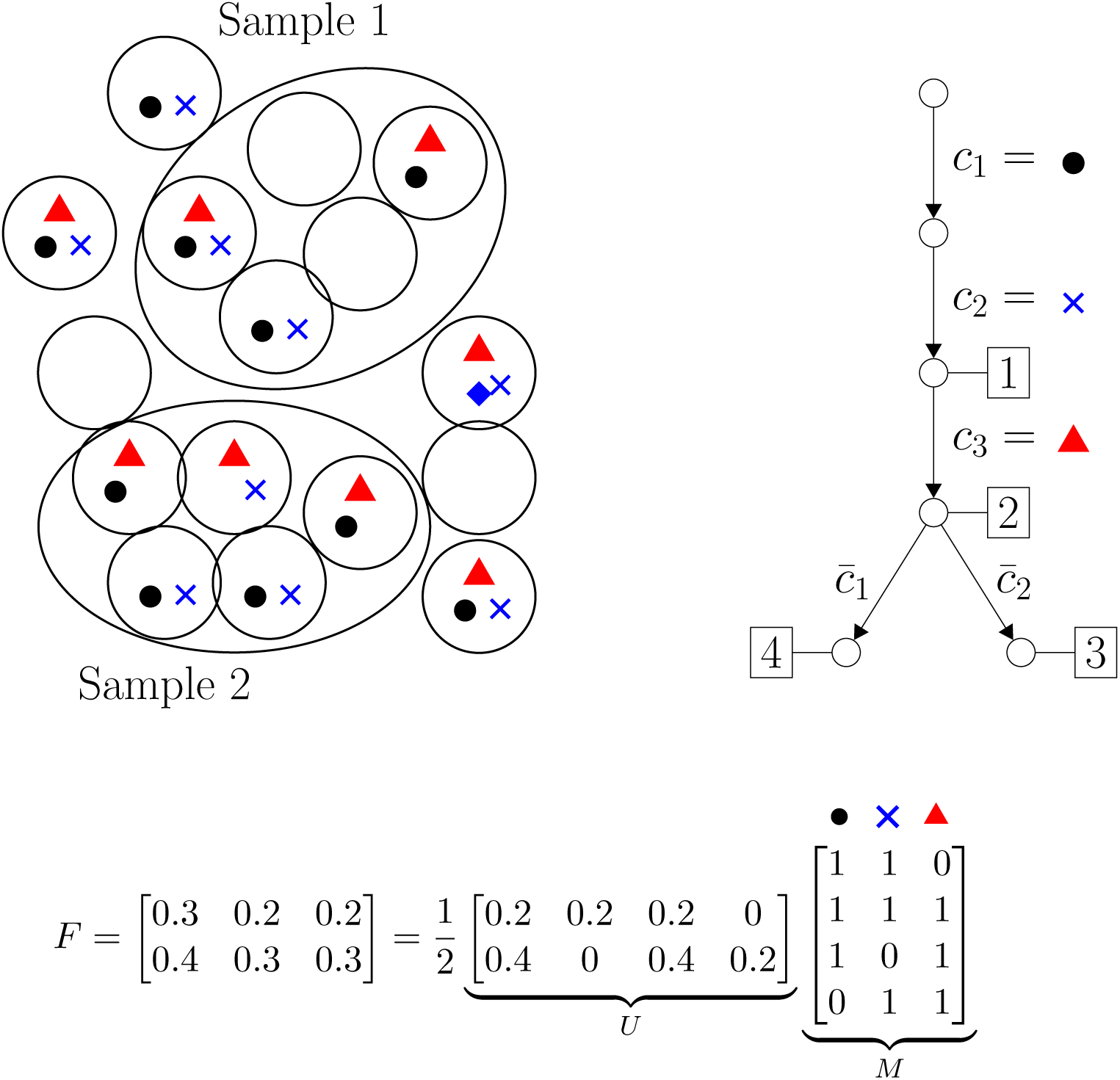
Example of the phylogenetic clonal reconstruction problem. On the left, the unknown clonal sub-populations of clones (top) and the the resulting VAF matrix (bottom). On the right, a solution for the Dollo(1)-VAFFP expressed as the product of matrices *U* and *M* (bottom) and a possible evolutionary history for the clones (top). Each colored dot represents a mutation.

The matrix factorization problem is simple when the clonal matrix *M* is known. In fact, once we have computed a clonal matrix *M*, the problem of finding a composition of samples, i.e. a usage matrix *U*, compatible with *M* consists simply of finding a matrix *U* such that 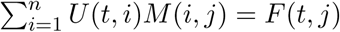 and 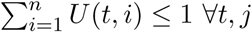.

Therefore, we decouple the 𝓟-VAFFP into two sub-problems: (1) the construction of the clonal matrix compatible with a phylogenetic model 𝓟, and (2) the search of the usage matrix which specifies the proportions of the proposed clones in the different samples.

The first of these problems is the main purpose of Section 3 in which we provide a ILP formulation for deciding if a clonal matrix admits a tree representation respecting a given phylogenetic model 𝓟. The second problem and the integration of both sub-problems is treated in Section 4.

### 2.2 The Incomplete Directed Perfect Philogeny Problem

The character-based phylogeny reconstruction problems we study in this paper are constrained versions of the general Incomplete Directed Perfect Phylogeny (IDP) [25]. In [25], the IDP problem asks for completing missing data in a binary matrix, where missing data are represented by the symbol ?, in such a way that the completed matrix is explained by a perfect phylogeny. More precisely, the input data is an *n* × *m* matrix *M*_?_, where *M*_?_(*i*, *j*) ∊ {0,1, ?} represents the absence, presence or uncertainty of a character *j* in the species *i* respectively. If a solution exists, then it consists of changing each ? into 0 or 1 obtaining a new binary matrix *M_s_* that has a directed perfect phylogeny.

A well known characterization of perfect phylogenies states that a binary matrix *M_s_* has a directed perfect phylogeny if and only if it has no *conflicting* pair of columns, which are two columns containing all the three configurations (0,1), (1,0), (1,1) — inducing the so called forbidden matrix. The problem of determining if a binary matrix has a perfect phylogeny, and to compute such perfect phylogeny if possible, has a linear-time algorithm [14,15]. Interestingly, the IDP problem has an efficient solution given by an *O*(*mn* log^2^(*m*+*n*))-time algorithm [25] when the phylogeny is directed, that is the root is known (it is the all 0s vector), otherwise, the problem of deciding whether there exists an unrooted solution of the incomplete input matrix is NP-complete [29]. There exists an ILP formulation for variants of the IDP problem, where the main question is to complete missing data in an input matrix on {0,1, ?} with the goal of minimizing the conflicting pairs [17]. Since finding a perfect phylogeny is easy, the main difficulty in solving the IDP problem consists of replacing each ? with a 0 or a 1 to minimize the number of conflicting pairs of columns.

### 2.3 ILP formulation for the IDP

In this section we revisit the ILP formulation proposed by Gusfield [17] for the IDP problem. The input of the problem is an incomplete *n* × *m* matrix *M*_?_. The goal is to decide if there exists a completion of the unknown entries of *M*_?_ resulting in a (complete) matrix admitting a Perfect Phylogeny. The main strategy of this approach is the minimization of the conflicts between pairs of characters. More precisely, in virtue of the Perfect Phylogeny Theorem, the IDP problem will have a solution if and only if the value of the problem is zero.

#### 2.3.1 Variables

We define a binary variable *Y* (*i*, *j*) for each unknown position of *M*_?_. With abuse of notation, *Y* (*i*, *j*) will be a constant for every known position of the matrix of value *M*_?_(*i*, *j*). Since the objective is to determine if two columns are in conflict, for every pair of columns *p*, *q* we define a binary variable *C*(*p*, *q*) that indicates the existence of a conflict between these two columns. To establish if two columns are in conflict, binary variables *B*(*p*, *q*, *a*, *b*) are defined for each pair of columns (*p*, *q*) and for each possible pair of values (*a*, *b*) ∊ {0,1}^2^. The variable *B*(*p*, *q*, *a*, *b*) indicates if for the (ordered) pair of columns (*p*, *q*) there exists a row *i* where *Y*(*i*,*p*) = *a* and *Y*(*i*,*q*) = *b*. Just as for the variable *Y* (*i*, *j*), if there exists a row of the matrix such that *Y* (*i*,*p*) = *a* and *Y* (*j*, *q*) = *b*, then *B*(*p*, *q*, *a*, *b*) = 1.

#### 2.3.2 Inequalities

For every pair of columns (*p*, *q*), every binary pair (*a*, *b*) ∊ {(1, 0), (0,1), (1,1)} and for every species *i*, the following set of inequalities

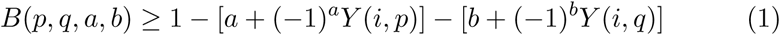

force the variable *B*(*p*, *q*, *a*, *b*) to be 1 if and only if the columns *p*, *q* exhibit the pair (*a*, *b*) in some row *i*. On the other hand, the following set of inequalities forces variables *C*(*p*, *q*) to be 1 when characters *p* and *q* are in conflict.

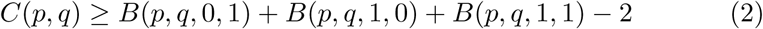

Since we are mainly interested in feasible solutions with no conflicts, we will consider the following alternative form of the previous constraint:

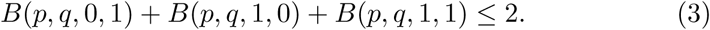

#### 2.3.3 Objective Function

Since we aim to minimize the number of conflicts, the objective function is defined as min ∑_(*P*,*q*)_ *C* (*p*, *q*).

By the previous discussion it is possible to state the problem of finding a completion with the minimal number of conflicts by considering the solution of the following minimization problem [17]: min ∑_(*p*,*q*)_ *C*(*p*,*q*) s.t. (1), (2). We stress the fact that decision problem of determine if an incomplete matrix admits a Perfect Phylogeny can be seen as checking if the former problem has zero value or equivalently a feasible solution for restrictions (1) and (3). The total number of variables and constraints in the formulation are in *O*(*nm* + *m*^2^) and *O*(*nm*^2^) respectively.

### 2.4 The Persistent Perfect Phylogeny and the IDP

Our strategy is based on the approach discussed by Gusfield [16] for the Persistent Phylogeny Problem, that is to decide if a binary matrix has a phylogeny representation for the Persistent model. The formulation proposed in [16] is based on two main properties:

1. Any instance *M* of the Persistent Phylogeny Problem can be reduced to an instance of an equivalent Incomplete Directed Perfect Phylogeny Problem on a matrix *M_e_*, called *extended matrix*, with some additional constraints [1], that is *M* has a Persistent Phylogeny if and only if *M_e_* has a perfect phylogeny.
2. The Incomplete Directed Phylogeny problem can be stated as an ILP problem by minimizing the number of conflicts between characters [17] according to the formulation presented in Section 2.3.

In the next section we extend this approach by generalizing the result presented in [1] in two different ways: First we extends the construction of the extended matrix to Dollo(*k*) and Camin-Sokal(*k*) models. Additionally, we generalize the construction to include the case in which the input matrix is incomplete in order to solve a more general problem: the Incomplete Directed Phylogeny Problem for the aforementioned phylogenetic models.

In the following we detail the construction proposed in [1] to reduce the Persistent Phylogeny Problem to an equivalent IDP instance. Given a (complete) binary matrix *M*, they propose an IDP problem on an (incomplete) extended matrix *M_e_* where each entry *M*(*i*,*j*) is replaced by two entries*M_e_*(*i*, *j*^+^) and *M_e_*(*i*, *j*^−^) as follows: if *M*(*i*, *j*) = 1 then *M_e_*(*i*, *j*^+^) = 1 and *M_e_*(*i*, *j*^−^) = 0; if *M*(*i*, *j*) = 0 then *M_e_*(*i*, *j*^+^) = *M_e_*(*i*, *j*^−^) =?. Given the input matrix *M_e_*, then a solution of Persistent Perfect Phylogeny is a binary matrix *M_s_* obtained by completing the entries of *M_e_* under the constraint that, for each pair (*M_e_*(*i*, *j*^+^),*M_e_*(*i*, *j*^−^)) of ? entries, the corresponding entries in the matrix *M_s_* must be the same, that is *M_e_*(*i*, *j*^+^) = *M_e_*(*i*, *j*^−^). Intuitively, the matrix *M_e_* corresponds to duplicate each column *j* corresponding to a character c into two columns *j*^+^, *j*^−^ corresponding to characters *c*^+^,*c*^−^, being *c*^+^ the gain of character c during evolution and *c*^−^ the loss of character *c*, in case *c* is a persistent character. Clearly, an entry *M*(*i*, *j*) = 1 means that the character *c* cannot be persistent. Thus the row *i* of *M_s_* is such that *c*^+^ is 1 and *c*^−^ is 0. Differently, an entry *M*(*i*, *j*) = 0 can be explained into two ways, either with the persistency of *c*, that is row *i* posses both characters *c*^+^ and *c*^−^ (both of them have values 1 in row *i*) or *c* does not occur in species row *i*, meaning that row *i* does not have characters *c*^+^ and *c*^−^ (both of them have values 0 in row *i*). Therefore:

#### Definition 2

([16]). Given an incomplete binary matrix *M_e_* and a set 𝓡 = {*R_i_*(*M_e_*) ≤ 0}_*i*∊[1,*r*]_ of *r* constraints on the entries of *M_e_*, the *Modified Incomplete Directed Perfect Phylogeny Problem for the set* 𝓡, denoted by MIDPP(*M_e_*,𝓡), asks to find, if it exists, a completion of matrix *M_e_* which admits a Perfect Phylogeny and satisfies all constraints in 𝓡

The fact that the obtained IDP includes some additional constraints makes more difficult to adapt the algorithm proposed in [25]. Therefore, we rather follow the approach proposed by Gusfield in [16] in which the restricted IDP is formulated as an ILP. Is easy to see that if every constraint in 𝓡 can be expressed as a linear constraint in terms of the matrix entries, then the problem MIDPP(*M*, 𝓡) admits a ILP formulation. The formulation can be obtained by simply adding the set of linear constraints 𝓡 to the former ILP formulation presented in Section 2.3.

## 3 The 𝓟 Incomplete Directed Phylogeny Problem

In this section we develop an ILP formulation for the following problem:

### Definition 3

(𝓟 Incomplete Directed Phylogeny Problem). Given a character-based phylogeny model 𝓟 and a incomplete binary matrix *M*, the 𝓟 Incomplete Directed Phylogeny Problem, denoted by 𝓟-IDP, asks for a completion *M_c_* of *M*, such that *M_c_* admits a phylogeny *T* under the model 𝓟, if such a completion exists.

Notice that if all entries of the input matrix *M* are known, then the problem corresponds to decide if *M* admits a phylogeny under the model 𝓟. In this paper we focus on Dollo(*k*) and Camin-Sokal(*k*). As we have already mentioned, we proceed by reducing the 𝓟-IDP on an instance *M* to an equivalent MIDPP(*M_e_*,𝓡*_M_*) instance where *M_e_* is a related extended matrix and 𝓡*_m_* is a set of linear restrictions. The later problem can thus be restated as an ILP.

### 3.1 The Dollo(*k*)-IDP

#### 3.1.1 Extended Matrix and Constraints for Dollo(*k*)

Let *M* be a binary (incomplete) matrix with *n* rows (species) and *m* characters. The extended matrix *M_D(k)_* for the Dollo(*k*) model is defined as follows:

- *M_D(k)_* has *n* rows and *m* × (*k* + 1) columns, where each character *j* of matrix *M* is associated to *k* + 1 columns in *M_D(k)_* denoted by *j*^+^,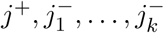.
- If *M*(*i*, *j*) = 1 then *M_D(k)_*(*i*, *j*^+^) = 1 and 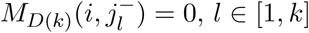.
- If*M*(*i*,*j*) = 0 or *M*(*i*,*j*) =? then *M_D(k)_*(*i*, *j*^+^) =? and 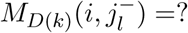 for each *l* ∊ [0; *k*].

For a character *j*, the column *j*^+^ represents the acquisition of character *j* while each of the 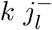 columns represents a possible loss of the gained character. In the case when *M*(*i*, *j*) = 1 then it is not possible for species *i* to lose the character *j* and the only possible configuration is *M_D(k)_*(*i*, *j*^+^) = 1 and 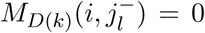, *l* ∊ [1,*k*]. Otherwise if *M*(*i*, *j*) = 0 then the character has either (1) never been acquired, or (2) been acquired, then lost along the path from the root to the species *i* of any solution. Therefore 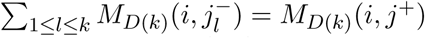.

Finally, if *M*(*i*, *j*) =?, that is the entry of *M* is missing, we must allow both the constraints for the case *M*(*i*, *j*) = 0 as well as *M*(*i*, *j*) = 1. consider both of the aforementioned relations, that is 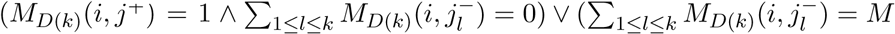. We capture both cases with with following relation between the entries of the extended matrix: 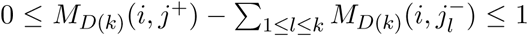. Our previous discussion leads to the following set of constraints for the matrix *M_D(k)_*:

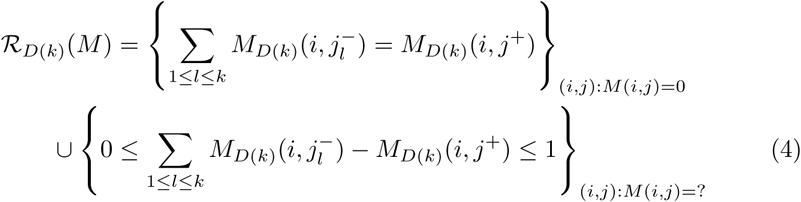

By an abuse of the notation it is possible to describe all restriction for the problem as:

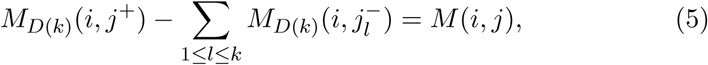

where the case *M*(*i*,*j*) =? is interpreted as 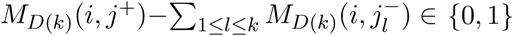.

When the context is clear, we will denote this set of restrictions as 𝓡*_D(k)_*. Figure 2 shows an example of the input matrix and its corresponding extended matrix.

**Figure 2:**
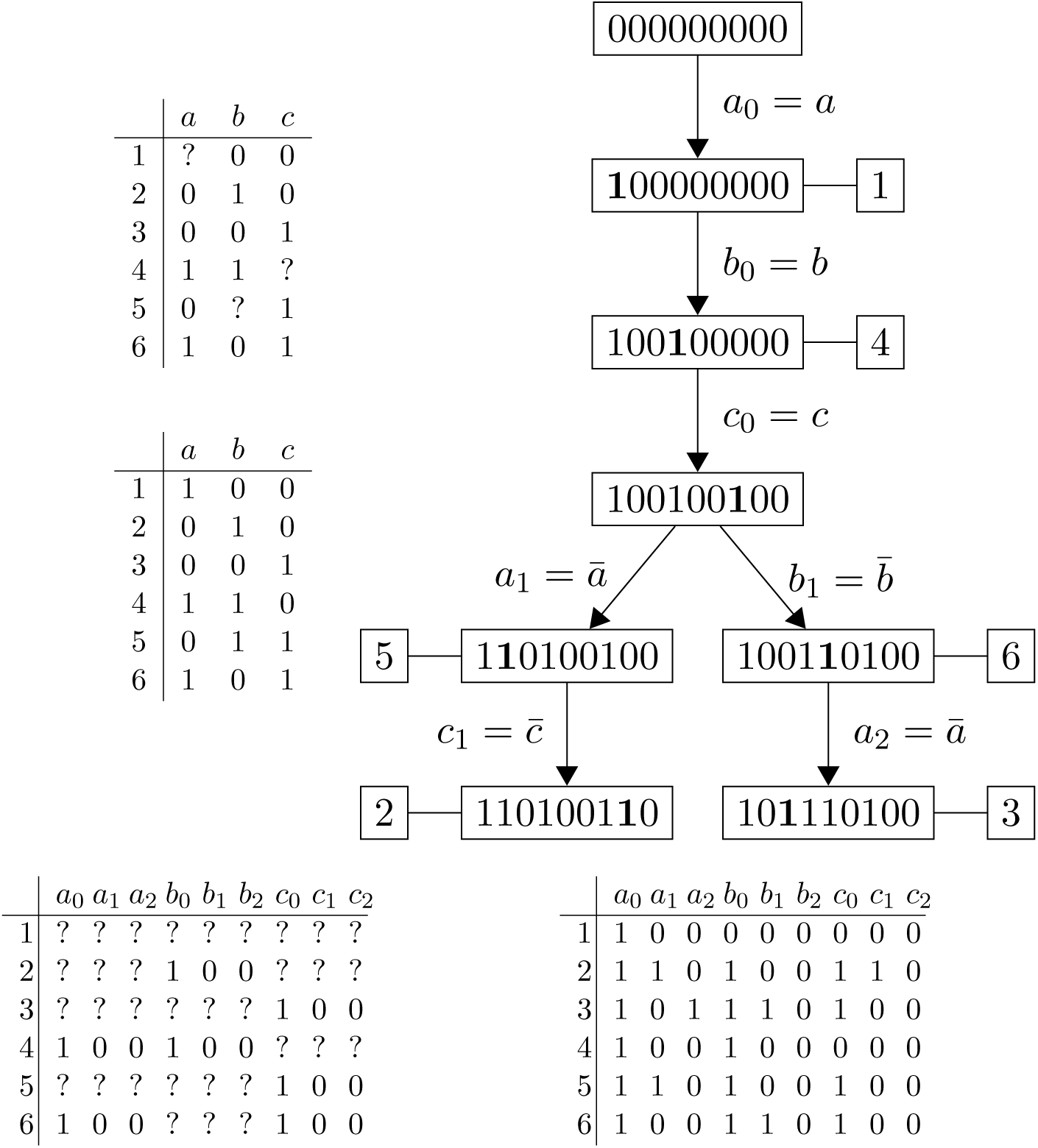
Input matrix *M* (top left), a Dollo(2) completion *M_c_* (center left) and its corresponding phylogeny tree *T* (top right). The *M_D_*_(2)_ extended matrix (bottom left) and a completion for the MIDPP(*M_D_*_(2)_, 𝓡*_D_*_(2)_) according to Theorem 4. In the tree, boldfaced character corresponds to changes between each node and its parent.

##### Theorem 4

*Let M be an incomplete binary matrix, and let MIDPP(M_D(k)_, 𝓡_D(k)_(M)) be the corresponding incomplete instance in the extended matrix M_D(k)_. Then there exist a completion M_c_ of M satisfying the Dollo(k) model if and only if MIDPP(M_D(k)_, 𝓡_D(k)_(M)) admits a solution.*

*Moreover, from any Dollo(k) completions M_c_ it is possible to obtain a solution of MIDPP(M_D(k)_, R_D(k)_(M)) and vice versa.*

##### Proof.

(⟹) Let *M_c_* be an completion for *M* that admits a Dollo(*k*) phylogeny *T_c_*. For each character *j* we relabel *T_c_* as follows: edges labeled *j*^−^ are relabeled from the set 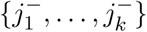 in such a way that no two edges receive the same label. Since *T_c_* is Dollo(*k*) phylogeny for *M_c_*, such a relabeling exists. Let *T*\* be the tree obtained from *T_c_* after relabeling. We denote by *M*\* the clonal matrix corresponding to *T*\*. Notice that *T*\* is a perfect phylogeny for *M*\*. Without loss of generality, we assume that *M*\* is a *n* × (*k* + 1) matrix: if a character is not present in *T*^*^ then we assign it a columns of zeroes in *M*^*^.

By our construction of *T*\* and *M*\*, if *M_c_*[*i*, *j*] = 1 then *M*\*[*i*, *j*^+^] = 1 and 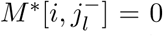 for all *l*, hence 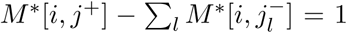. If *M_c_*[*i*, *j*] = 0 then either (1) *M*\*[*i*, *j*^+^] = 1 and exactly one of the entries 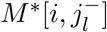 with 1 ≤ *l* ≤ *k* is equal to one, or (2) *M*\*[*i*, *j*^+^] = 0 and 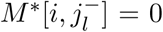 for all *l*. In both cases, 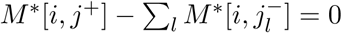.

Since *M_c_*[*i*, *j*] = 1 ⟹ *M*\*[*i*, *j*] ∊ {1, ?} and *M_c_*[*i*, *j*] = 0 ⇒ *M*\*[*i*, *j*]∊{0, ?}, the above argument implies that the variables corresponding to entries of *M*\* satisfy the constraints in 𝓡*_D(k)_*.

(⇐) Conversely, let *M* be an incomplete binary matrix and let *M*\* be a solution of MIDPP(*M_D(k)_*, 𝓡*_D(k)_*). We will proof that *M* has a completion *M_c_* with a Dollo(*k*) phylogeny *T_c_*.

Let *T*\* be the perfect phylogeny tree of *M_c_*. We construct the phylogeny tree *T_c_* from *T*\* by replacing each label 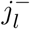 with *j*^−^ respectively. Since the matrix *M*\* satisfies restrictions in (5) then 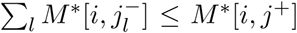, thus column *j*^+^ is bigger (component-wise) than all columns 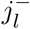. Hence, in the tree *T_c_* the edge *j*^+^ is in the path to the root from any edge labeled with *j*^−^. We conclude that the tree *T_c_* is a Dollo(*k*) phylogeny and we denote by *M_c_* its corresponding binary matrix.

By our construction of *T_c_*, 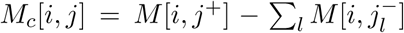 for each known entry of a species *i* and character *j*. Hence *M_c_* is a completion of *M*.

### 3.2 The Camin-Sokal(*k*) IDP

#### 3.2.1 Extended Matrix and constraints for Camin-Sokal(*k*)

Let *M* be a incomplete binary matrix with *n* species and *m* characters. The extended matrix *M_CS(k)_* for the Camin-Sokal(*k*) model is defined as follows:

- *M_CS(k)_* has *n* rows and *m* × *k* columns; each character *j* of matrix *M* is associated to *k* columns in *M_CS(k)_* denoted by 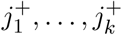.
- If *M*(*i*, *j*) = 0 then 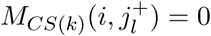, ∀*l*.
- If *M*(*i*, *j*) = 1 or *M*(*i*, *j*) =? then 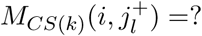,*l* ∊ [1,*k*].

Every group of columns 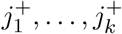 represent the possible gain of character *j* in the resulting phylogenetic tree. In every feasible solution, a character can be gained at most once on any path from the root to a leaf, therefore we define the set following set of constrains for the extended matrix *M_CS(k)_*:

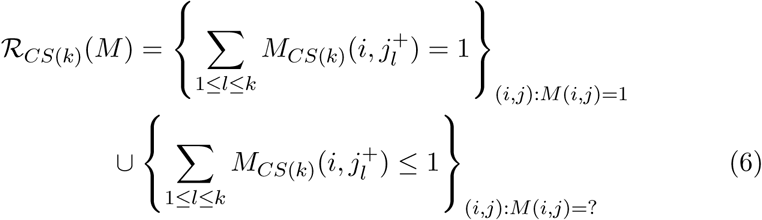

Similarly to the Dollo(*k*) case, we can express the restriction set as:

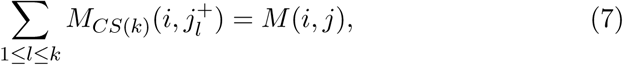

for the case *M*(*i*, *j*) =? the equation is interpreted as 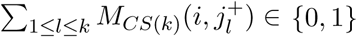.

##### Theorem 5

*Let M be an incomplete binary matrix, and let MIDPP(M_CS(k)_, R_CS(k)_ (M)) be the corresponding incomplete instance in the extended matrix M_CS(k)_. Then there exist a completion M_c_ of M satisfying the Camin-Sokal(k) model if and only if MIDPP(M_CS(k)_, R_CS(k)_ (M)) admits a solution.*

*Moreover, from any Camin(k) completions M_c_ it is possible to obtain a solution of MIDPP(M_CS(k)_, R_CS(k)_(M)) and vice versa.*

##### Proof

(⇒)Let *M_c_* be a completion of *M* admitting a Camin-Sokal(*k*) phylogeny *T*. We relabel the edges of *T* assigning to each edge labeled as *j*^+^ a different label in the set 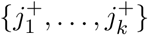. Let *T*\* be the tree obtained from *T* after relabeling, it is easy to see that *T*\* represent a perfect phylogeny for the new species set. We denote by *M*\* the clonal matrix corresponding to *T*\*. Morever, we assume that *M*\* has all colums associated to the characters in the set 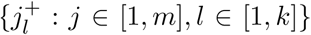. Otherwise, in the case that a character is not present in the tree then we fill its corresponding columns on *M*\* with zeros. Since in the phylogeny *T* a species *i* never gains a character *j* that it does not possess, then 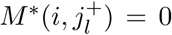 for *l* ∊ [1,*k*]. Thus, matrix *M*\* is a completion of the extended matrix *M_D(k)_*. Let verify that *M*\* entries satisfies the restrictions in 𝓡_*CS*(*k*)_. Since *T*\* is perfect phylogeny tree, then it holds that 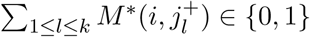 for all species *i* and character *j*. Additionally, for each species *i* containing a character *j*, the path from *i* to the root contains only one edge labeled with *j*^+^ meaning that 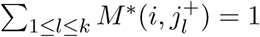.

(⇐) Let *M*\* be a solution of MIDPP(*M_D(k)_*, 𝓡*_D(k)_*) for an input matrix *M*. Let *T*\* be the perfect phylogeny tree of *M*\*. We construct the tree *T* from *T*\* by relabeling all edges with label 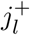,*1* ∊ [1,*k*] with *j*^+^. Since *T*^*^ represents a perfect phylogeny, then in each path from the root to an species no label is duplicated. Therefore the tree *T* represents a phylogeny respecting the Camin-Sokal(*k*) model. We denote by *M_c_* the clonal matrix corresponding to *T*. Since *M*\* satisfies (7) we conclude that the matrix *M_c_* is a completion of *M*.

An instance of the previous construction is shown in Figure 3 of the Camin-Sokal(2) Phylogeny for the input matrix in Figure 2. Finally, we can state the Camin-Sokal(*k*) Phylogeny Reconstruction Problem as the minimization problem min ∑*_(pq)_ C*(*p*,*q*) s.t. (1), (2) and (6), or a feasible solution of the restriction set (1), (3) and (6).

**Figure 3:**
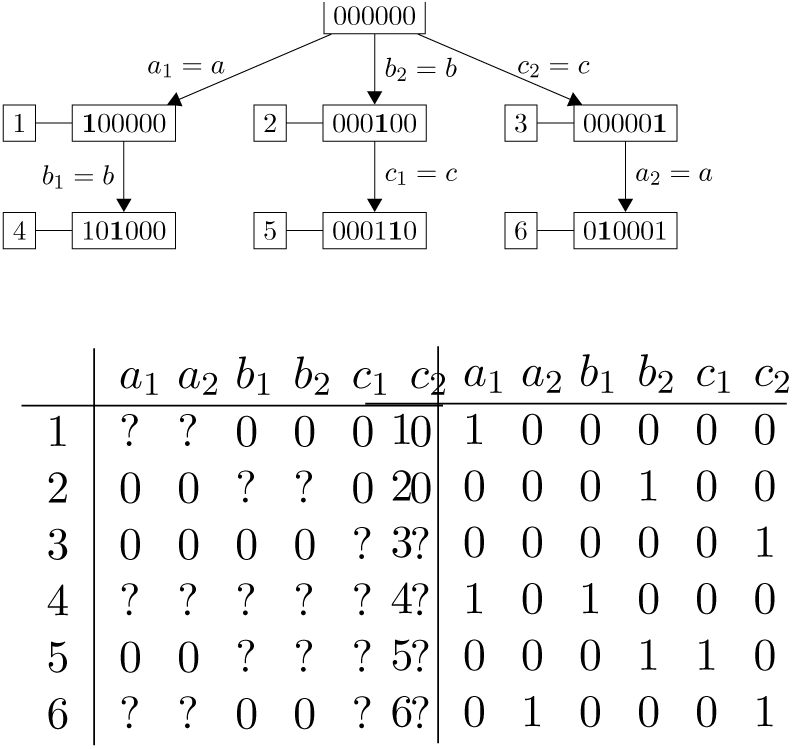
A Camin-Sokal(2) phylogeny (top) for the input matrix M and its completion of Figure 2. In the bottom left we represent the extended matrix, while in the bottom right we represent the corresponding completion for the MIDPP(*M_CS_*(_2_),*R_CS_*(_2_)) according to Theorem 5. In the tree, boldfaced character corresponds to changes between each node and its parent.

## 4 The Clonal Reconstruction Problem

While Section 3 focuses on the Incomplete Phylogeny Problems where the instance is an incomplete binary matrix, this section is dedicated to tumoral multisample instances. More precisely, in this section we present an ILP formulation for the 𝓟-VAFFP. Let us recall that a 𝓟-VAFFP instance is a *p* × *m* frequency matrix *F*, and a number of clones *n*. The goal is to find two matrices *U* and *M*, respectively the *p* × *n usage* matrix, representing the composition of the samples in terms of clones, and the *n* × *m clonal* matrix, representing the desired tree. Moreover, *M* represents a phylogeny satisfying the rule 𝓟, and the expression 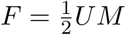, which guarantees that the frequency of the leaves of the tree are actually equal to those in the input matrix *F*.

In our approach, the most fundamental variables are those corresponding to usage, clonal and extended matrices, which we denote by *U*, *M* and *M_e_* respectively. The extended matrix is constructed according to Section 2 and following the phylogeny model 𝓟. More precisely, we will have entries *U*(*t*, *i*), *M*(*i*, *j*), *M_e_*(*i*, *j_l_*), for each sample *t* ∊ [1,*p*], clone *i* ∊ [1,*n*] and mutation *j* ∊ [*i*, *m*].

First, we guarantee that each row of the usage matrix *U* is actually the composition of the sample, where the entry *U*[*t*, *i*] is the fraction of cells in the sample *t* that belong to the clone *i*, by imposing

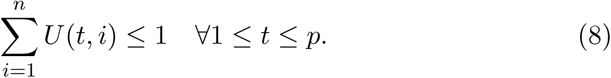

Then the constraints on the matrices *M* and *M_e_* are those of Section 2.3 and guarantee that *M* encodes a phylogeny *T* whose characters are the input mutations and *T* is consistent with the model 𝓟.

On the other hand, we must guarantee that the clonal matrix *M* admits a phylogeny under the 𝓟 model. As it was discussed in Section 3, it is possible to state the 𝓟 phylogeny reconstruction problem as a solution for an IDP problem on the corresponding extended matrix (Theorem 4 and Theorem 5).

The relation between the matrices *F*, *U*, and *M*, as stated in the equation 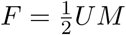, is enforced by the set of constraints

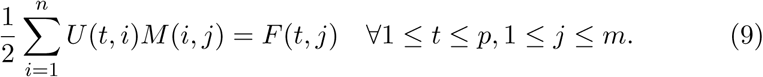

Unfortunately, Equation 9 gives a set of quadratic constraints that cannot be solved directly via ILP. Therefore, we need to replace those constraints with the following linear constraints that need the set of auxiliary binary variables *X*(*t*, *i*, *j*) where, as usual, 1 ≤ *t* ≤ *p*, 1 ≤ *i* ≤ *n*, 1 ≤ *j* ≤ *m*. More precisely, each variable *X*(*t*, *i*, *j*) is equal to the product *U*(*t*, *i*)*M*(*i*, *j*).

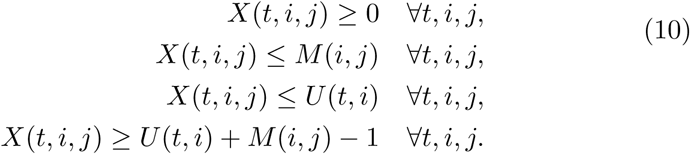

Thus, the 𝓟-VAFFP corresponds to finding a feasible solution with the linear constraints (1), (3), (8), (10), (5) for the Dollo(*k*) model, and (1), (3), (8), (10), (7) for the Camin-Sokal(*k*) model.

Finally, since the matrix *M_e_* has at most *km* columns, our complete formulations has *O*(*nkm* + *k*^2^*m*^2^ + *mpn*) variables and *O*(*k*^2^*m*^2^ + *npm*) constraints.

### 4.1 Clonal Reconstruction admitting errors

Since the frequency matrix *F* is obtained experimentally, via mapping reads to the reference genome, the measured frequency is only an approximation of the actual frequency. For this reason, we extend our formulations to incorporate frequency errors and we pick the minimization of the overall errors as our objective function.

More precisely, we introduce the set of variables *E*(*t*, *j*) that represent the error in the measure of the input frequency *F*(*t*, *j*). Notice that 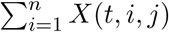 is (implicitly) our estimated frequency, therefore the following constraints determine the value of the variables *E*(*t*, *j*) as the difference between the input frequency and the estimated frequency.

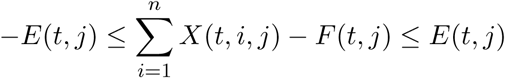

Since now our goal is to minimize the overall error introduced in the reconstruction, the objective function is:

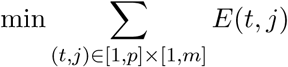

### 4.2 gppf

We implemented our approach with a Python program called `gppf` that receives a frequency matrix *F* and the evolution model (Persistent, Dollo(*k*), Camin-Sokal(*k*)). The program computes the corresponding ILP which is fed to Gurobi 6.5.2. Moreover, our program receives the solution computed by Gurobi and returns a tree, provided that Gurobi has been able to find a feasible solution. The program `gppf` is available at https://github.com/AlgoLab/gppf.

The parameters of the implementation are the maximum number of clones that a solution can use (expressed as the percentage of the number of mutations), the maximum time permitted for each execution, and the parameter *k* associated to the model Dollo(*k*) and the Camin-Sokal(*k*) in the formulation. Moreover, we have introduced a timeout on the running time, since the generated ILP problem is often large and its resolution could require a considerable amount of time. We exploit the fact that Gurobi can be halted at any time and it returns the best feasible solution computed so far. Hence, imposing a timeout allows the ILP solver to compute a solution with a small total error.

## 5 Experimental Results

All experiments have been performed on an Ubuntu 14.04 server with four 8-core Intel Xeon E5-4610v2 2.30GHz CPUs (hyperthreading was enabled for a total of 16 threads per processor).

The goals of the experimental analysis have been two: to test the hypothesis that evolution models that do not satisfy the infinite site assumption can actually provide better predictions, and to assess the computational feasibility of our approach. More precisely, besides the Perfect Phylogeny model, we have tested the Persistent Phylogeny, the Dollo(*k*), and the Camin-Sokal(*k*) models on both simulated and real data. The size of the instances are typical for real data applications such as liquid cancer and in particular Leukemia.

We have simulated some datasets — more precisely, frequency matrices *F* — according to the following steps:

1. we have generated a clonal *n* × *m* matrix *M* with the simulation tool `ms` [19], obtaining a Perfect Phylogeny on *n* clones and *m* mutations.
2. We have flipped at most 30% of the 0s of *M* into 1s, uniformly at random. This allows us to have phylogenies that are not necessarily perfect.
3. We have generated a *p* ×*n* usage matrix *U* assigning to each clone a proportion in each sample. The frequencies are chosen randomly following a Dirichlet distribution.
4. We have multiplied *U* and *M* to generate a *p* × *m* frequency matrix *F*.

Each generated matrix is given as input to `gppf` with different evolution models. We remark that we do not compare the predicted clonal matrix *M* with the original, since different models can generate diverse clonal evolution trees.

We evaluate the computed solutions according to the following measure which is a ratio where smaller values correspond to better predictions:

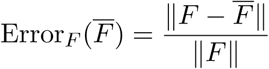

Where *F* is the input frequency matrix, *F*̅ is the frequency matrix inferred by the solution, and ‖*A*‖= [∑*_ij_*|*a_ij_*|^2^]^1/2^ is the Frobenius norm. This metric give us the ratio between the total error and the optimal value, therefore it is not too dependent on the actual values.

Previous works focused on Perfect Phylogeny as the evolutionary model, thereby restricting the attention to a number of clones equal to the number of mutations. More general evolutionary models do not have this constraint, that is the number of clones might be different. We have investigated the effect of choosing different values of the maximum number of clones. More precisely, we have considered the number of clones to be at most 100%, 80%, 60% and 40% of the number of clones in the instance. Notice that the actual number of clones in the actual solution might be smaller.

### 5.1 Simulated Data

For the simulated data, we have generated two different datasets:

**Exp. 1** contains 100 frequency matrices composed of 6 samples and 10 mutations. Matrices are generated from a 20 × 20 clonal matrix *M*. The phylogenetic models tested in this set are: Perfect, Persistent, Dollo(2) and Camin-Sokal(2).
**Exp. 2** contains 30 frequency matrices with 12 samples and 25 mutations, generated by a 25 × 50 clonal matrix *M*, The models tested in this set are: Perfect, Persistent, Dollo(2) and Dollo(4). Given the results of the previous experiment we decided to abandon the Camin-Sokal model and to evaluate different parameters for the Dollo model.

Figures 4 and 5 show how the error of each solution varies as a function of the running time for both experiments. We notice that the executions on the Perfect Phylogeny model quickly reaches a plateau, while the same is not true for the Persistent Phylogeny and the Dollo or Camin-Sokal models, where a longer time is needed. Moreover, the plots for the Dollo and Camin-Sokal models hint that the plateau is not actually reached. In fact, those models are more general than the Persistent Phylogeny, hence the optimumin those case should have an error that is at least as good as the one found for the Persistent Phylogeny model.

**Figure 4:**
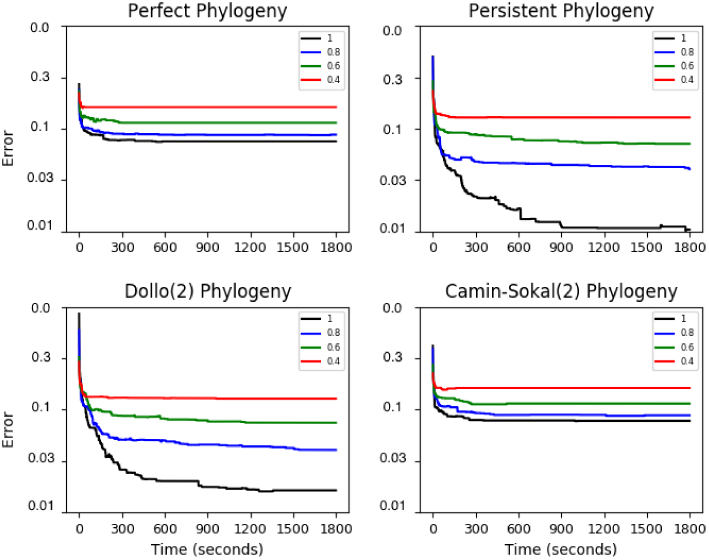
Average error of the solutions computed for Experiment 1 as a function of the running time. The error is on a logarithmic scale. The running time is in seconds.

**Figure 5:**
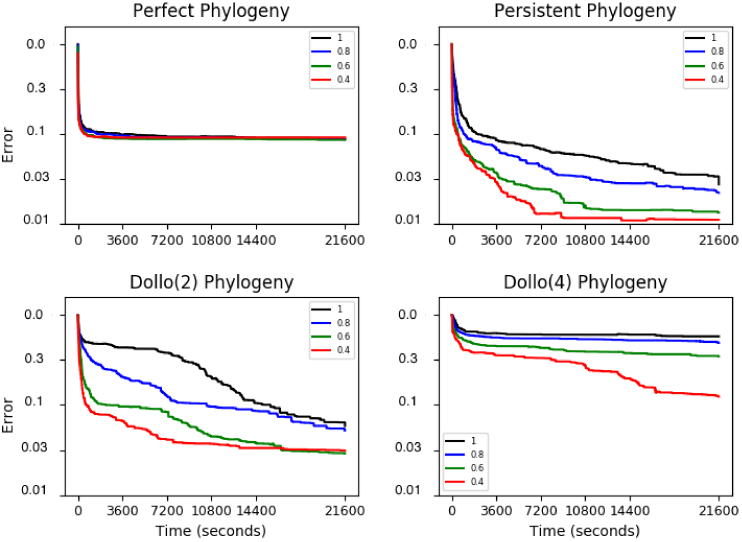
Mean error of the solution computed for Experiment 2 as a function of the running time. The error is on a logarithmic scale. The running time is in seconds.

Finally, the analysis of Figures 4 and 5 leads us to set a time limit for the running time equal to 5 minutes for Experiment 1 and 6 hours for Experiment 2, since allowing a large time limit results in only marginal improvements of the quality of the solutions computed.

Figures 6, 7 compare the total error of the solutions obtained under different phylogenetic models and different upper bounds on the number of clones for the Experiments 1 and 2. We recall that we have set a timeout of 5 minutes and 6 hours respectively for Experiment 1 and 2. Additionally, Tables 1 and 2 report the number of input instances where considering more general phylogeny models allows to compute solutions that are better than those conforming to the Perfect Phylogeny model.

**Figure 6:**
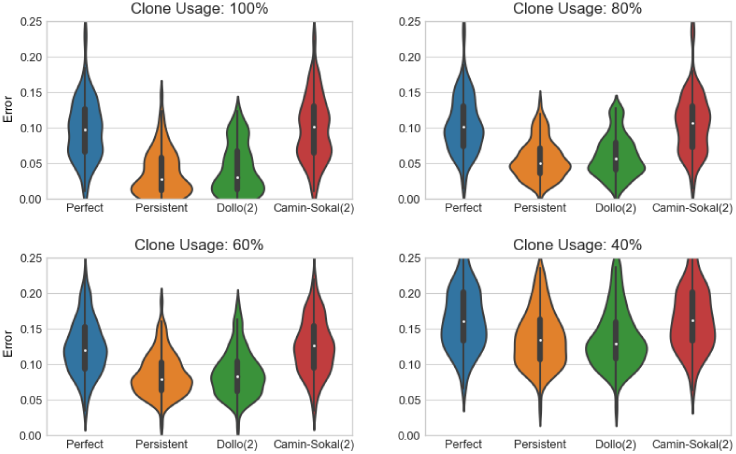
Comparing the error of the solutions for different evolution models: Experiment 1. The figure represents the distribution of the error for different values of the ratio between the clone limit (the maximum number of clones allowed) and the number of mutations. The Persistent Phylogeny and the Dollo(2) models consistently give the better results.

**Figure 7:**
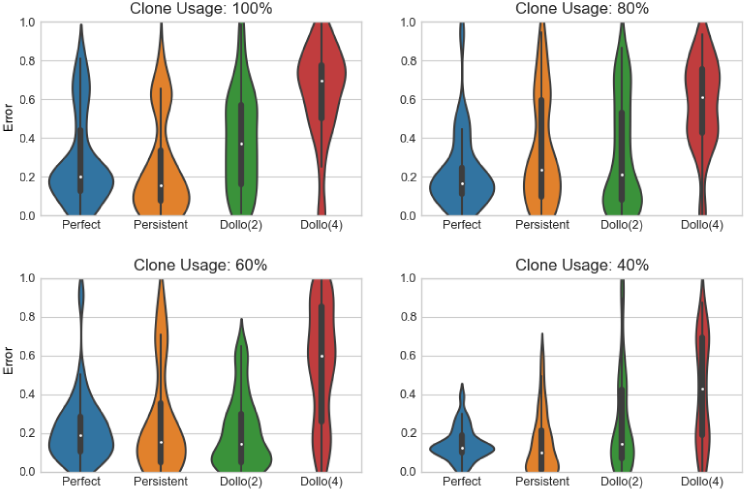
Comparing the error of the solutions for different evolution models: Experiment 2. The figure represents the distribution of the error for different values of the ratio between the clone limit (the maximum number of clones allowed) and the number of mutations.

**Table 1:** Comparison between evolution models on Exp. 1. Each entry contains the number of instances (out of the 100 instances with same ratio between the maximum number of clones and the number of mutations) where the formulations based on the Persistent Phylogeny, Dollo(2), Camin-Sokal(2) models obtain a total error that is smaller than a certain fraction of the one obtained with the Perfect Phylogeny model.

| Ratio<br># clones /<br># mutations | Ratio total er-<br>ror / total error<br>on perfect phy-<br>logeny | Persistent | Dollo(2) | Dollo(4) |
| --- | --- | --- | --- | --- |
| 1 | $\leq 100\%$ | 100/100 | 94/100 | 47/100 |
| | $\leq 90\%$ | 99/100 | 92/100 | 18/100 |
| | $\leq 80\%$ | 97/100 | 86/100 | 10/100 |
| | $\leq 50\%$ | 74/100 | 66/100 | 0/100 |
| 0.8 | $\leq 100\%$ | 100/100 | 97/100 | 53/100 |
| | $\leq 90\%$ | 99/100 | 91/100 | 11/100 |
| | $\leq 80\%$ | 89/100 | 84/100 | 4/100 |
| | $\leq 50\%$ | 44/100 | 39/100 | 0/100 |
| 0.6 | $\leq 100\%$ | 98/100 | 92/100 | 51/100 |
| | $\leq 90\%$ | 91/100 | 83/100 | 2/100 |
| | $\leq 80\%$ | 74/100 | 62/100 | 1/100 |
| | $\leq 50\%$ | 13/100 | 17/100 | 0/100 |
| 0.4 | $\leq 100\%$ | 90/100 | 93/100 | 79/100 |
| | $\leq 90\%$ | 70/100 | 74/100 | 0/100 |
| | $\leq 80\%$ | 43/100 | 43/100 | 0/100 |
| | $\leq 50\%$ | 0/100 | 0/100 | 0/100 |

**Table 2:** Comparison between evolution models on Exp. 2. Each entry contains the number of instances (out of the 30 instances with same ratio between the maximum number of clones and the number of mutations) where the formulations based on the Persistent Phylogeny, Dollo(4), Camin-Sokal(4) models obtain a total error that is smaller than a certaun fraction if the error obtained with the Perfect Phylogeny model.

| Ratio<br># clones /<br># mutations | Ratio total er-<br>ror / total error<br>on perfect phy-<br>logeny | Persistent | Dollo(2) | Dollo(4) |
| --- | --- | --- | --- | --- |
| 1 | $\leq 100\%$ | 19/30 | 09/30 | 07/30 |
| | $\leq 90\%$ | 18/30 | 08/30 | 05/30 |
| | $\leq 80\%$ | 14/30 | 07/30 | 04/30 |
| | $\leq 50\%$ | 11/30 | 05/30 | 02/30 |
| 0.8 | $\leq 100\%$ | 12/30 | 13/30 | 02/30 |
| | $\leq 90\%$ | 11/30 | 13/30 | 02/30 |
| | $\leq 80\%$ | 11/30 | 13/30 | 02/30 |
| | $\leq 50\%$ | 09/30 | 07/30 | 02/30 |
| 0.6 | $\leq 100\%$ | 16/30 | 19/30 | 05/30 |
| | $\leq 90\%$ | 15/30 | 18/30 | 05/30 |
| | $\leq 80\%$ | 15/30 | 16/30 | 05/30 |
| | $\leq 50\%$ | 08/30 | 11/30 | 01/30 |
| 0.4 | $\leq 100\%$ | 18/30 | 13/30 | 06/30 |
| | $\leq 90\%$ | 18/30 | 11/30 | 04/30 |
| | $\leq 80\%$ | 16/30 | 11/30 | 03/30 |
| | $\leq 50\%$ | 12/30 | 07/30 | 02/30 |

W.r.t. the number of allowed clones, the more general models result in better predictions, as expected. There is a similar trend when comparing different evolution models, that is the Perfect Phylogeny model is usually outperformed by the Persistent Phylogeny and the Dollo(2) models. In this case, the much larger search space of more general models does not allow the ILP solver to find a near-optimal solution. Still, Tables 1, 2 show that, in almost all instances, a general phylogeny model outperform the results of the Perfect Phylogeny solution.

Notice that Camin-Sokal (Figure 6) and Dollo(4) (Figure 7) are not able to match the quality of the predictions under the Perfect Phylogeny model. Nevertheless, we note that Persistent and Dollo(2) model obtain better results than the Perfect Phylogeny, especially when the allowed number of clones is small. The Persistent model obtains better results in more than half the simulations even with all, or almost all, the clones. Experiments 1 required 420 CPU hours, while Experiment 2 required 2880 CPU hours.

### 5.2 Real data: Chronic Lymphocytic Leukemia

To test the accuracy gppf on real cancer data, we run the ILP formulation on the dataset provided in [27]. We expect our tool to confirm the main findings of that paper. Whole-Genome Sequencing (WGS) was used to track subclonal heterogeneity in 3 chronic lymphocytic leukemia (CLL) patients subjected to repeated cycles of therapy. Between 14 and 22 mutations per sample were predicted to alter protein-coding sequencing. WGS analysis confirmed the presence of copy number aberrations (CNAs) in all patients. We point out that our method is unable to fully consider copy number aberrations, since our model only consider the presence of absence of a mutation. Consequently, these datasets are among the most difficult to manage with our approach. Still, we want to compare our predictions with those in the literature: we will show that we are able to confirm almost all the findings in the relevant studies.

The choice of the Chronic Lymphocytic Leukemia datasets was due mostly to the reduced number of somatic mutations in liquid tumors, that allowed us to calculate an optimal solution in a reasonable amount of time. Since the Persistent model seemed the most promising from the experimental results we decided to use this particular model to infer the mutational evolutionary history of the CLL dataset.

The inferred mutation lineage for CLL patient 077, which consists of 5 samples and 16 somatic mutations, is shown in Figure 8; the Persistent Phylogeny model was computed in approximately 3 days. The inferred tree is consistent with the clonal expansion proposed in the original study [27]. The driver mutation SAMHD1 is successfully inferred by `gppf` as well as the fact that there are 4 major lineages. Our prediction contains five leaves instead of the expected four leaves. The reason is that one of leaves in our tree is the result of the mutation loss of OCA2. We argue that such loss corresponds to the (only) CNA described in the original study.

**Figure 8:**
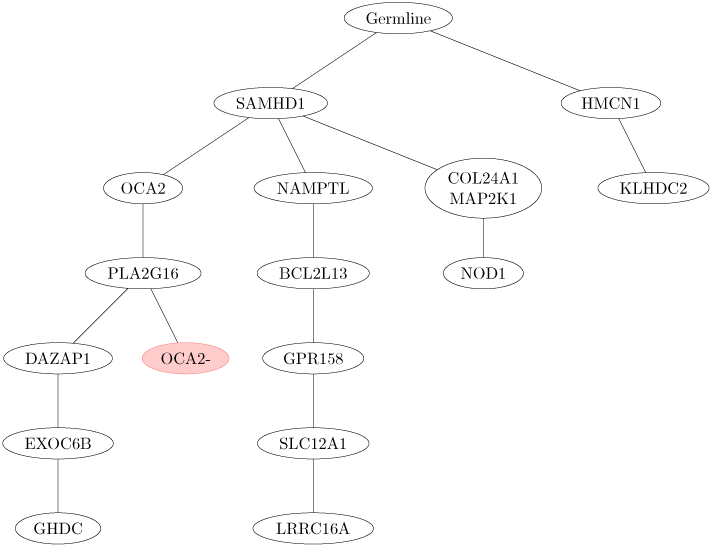
Solution computed by *gppf* for patient CLL077. Nodes with red background represent backmutations *i.e.*, mutation losses). Therefore we have one clonal expansion where mutation *OCA2* has been lost. The driver mutations of this tree are those of [27].

We cannot directly compare the persistent tree we inferred with AncesTree [9] (Figure 9), because the latter infers only seven of the 16 mutations present in the sample. In order to perform such comparison we had to restrict the instance to contain only the mutations that are also in the solution computed by AncesTree. The output is presented in Figure 11 and shares several structural similarities with the AncesTree solution. Moreover, we would also like to point out that our solution for the restricted instance has no errors (and is therefore optimal).

**Figure 9:**
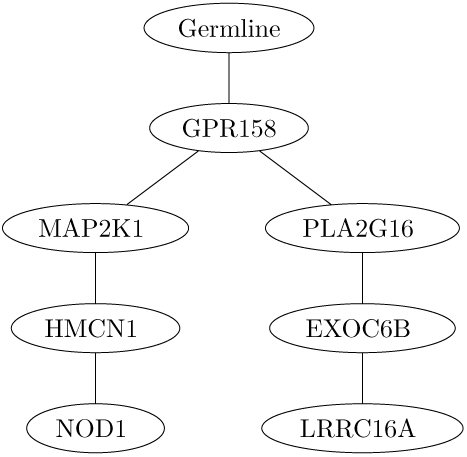
Solution computed by by AncesTree on CLL077.

We have compared our predictions with those of PhyloSub [20] (Figure 10) as well. PhyloSub clusters together some mutations in the same clone, while we infer a tree in which each mutation correspond to a vertex, except for mutations where, given their inferred mutation profile (*i.e.*, the presence of each mutation in the clone), `gppf` is unable to predict their ancestry relationship. The cluster detected by PhyloSub containg mutations OCA2, PLA2G16, DAZAP1, EXOC6B, HMCN1 and GHDC is preserved in the solution predicted by `gppf` that instead of clustering the mutations defines a lineage between them, with the exception of HMCN1 that is instead child of the germline. The same applies for the cluster that includes NAMPTL, BCL2L13, GRP158, SLC12A1 and SAMHD1, but the latter is identified as a driver by `gppf` for which the two previous cluster are children. The main difference is mutation LRRC16A that for PhyloSub and `gppf` is descendant of two different clusters. The last cluster identified by PhyloSub containing mutations COL24A1, MAP2K1, NOD1, HMCN1 and KlHDC2 is being separated by `gppf` which predicts that the first three mutations are indeed derived from the driver SAMHD1, while the last two mutated directly from the germline.

**Figure 10:**
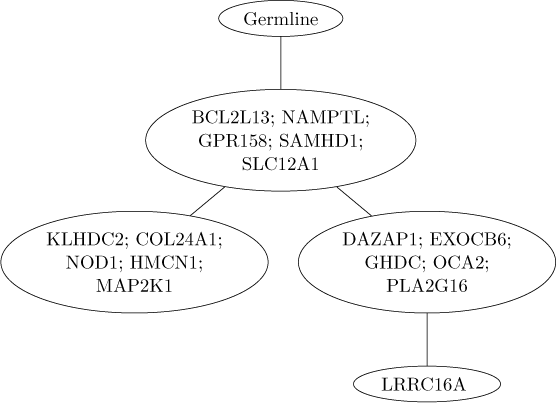
Solution computed by PhyloSub on CLL077.

**Figure 11:**
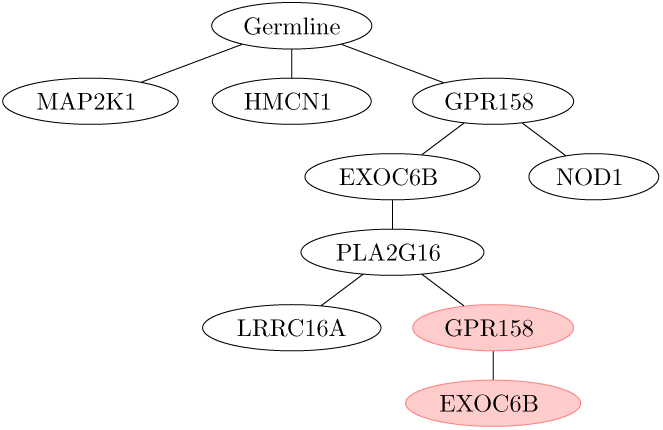
Solution computed by `gppf` under the Persistent Phylogeny model for the restricted instance solved in [9] with AncesTree. The solution computed by AncesTree on the same instance is in Figure 9.

Patient CLL006 presented a total of 10 somatic mutations in 5 samples. It is important to notice that patient CLL006 has been reported to have trisomy 12; this particular disease has been shown [7] to be associated with chromosome 14q deletions and therefore it is expected to report a large amount of back-mutations. Indeed, as shown in Figure 12, `gppf` identify a total of 7 mutations that have been lost in the cancer progression. Moreover the tumor progresses as a chain of mutations in its early stage as reported in the original study and our model correctly infer MED12 as a driver mutation.

**Figure 12:**
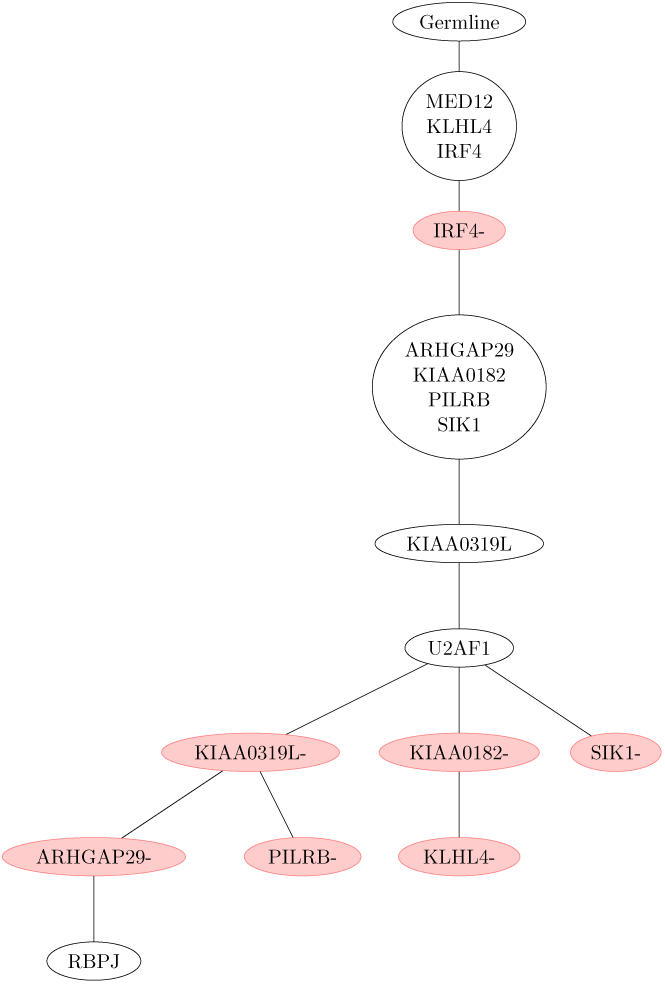
Solution computed by `gppf` for patient CLL006. Nodes with red background represent backmutations (*i.e.*, mutation losses). Given the presence of trisomy 12 in this patient a large amount of backmutations are expected in the progression of the tumor. In this case it is unknown the relative order of the mutations acquired in the same node of the tree. We confirm the driver mutation MED12 of [27].

The last patient in the study was CLL003 in which a total of 20 somatic mutations were found among 5 samples. While `gppf` (Figure 13) and the original study do not infer the same driver mutations, the overall cancer progression is very similar; both in fact report three main lineages and a significant loss of mutation in the last lineage. A total of 3 CNAs was reported in the study while `gppf` identifies 2 losses: CHRNB2 and NRG3. Still, a CNA can be a duplication, not necessarily a mutation loss.

**Figure 13:**
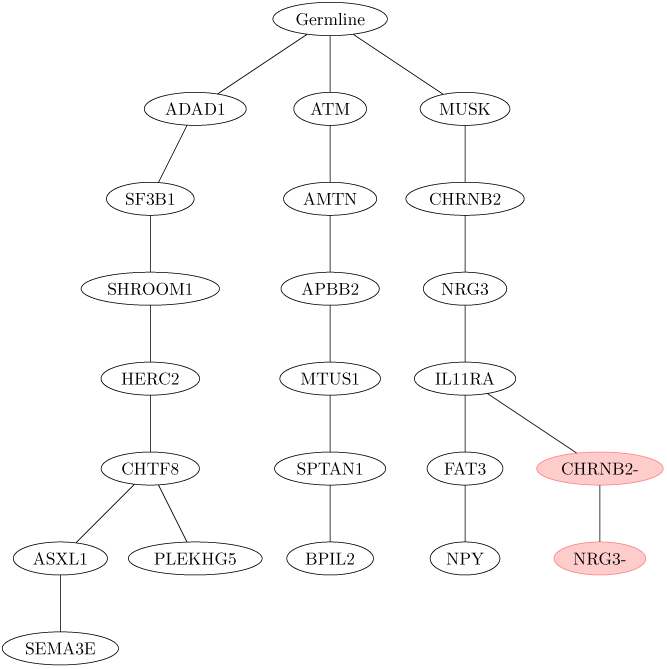
Solution computed by `gppf` for patient CLL003. Nodes with red background represent backmutations (*i.e.*, mutation losses). The overall tree topology of the cancer progression is similar to the one proposed in the original study, with minor differences in driver mutations.

## 6 Conclusions and Future Work

In this paper we have proposed a ILP formulation of the problem of reconstructing the evolutionary history of tumors, where the evolutionary tree is character-based and can violate the infinite site assumption of the Perfect Phylogeny model.

First, we have proposed an ILP framework for the Dollo(*k*) and Camin-Sokal(*k*) models — *k* is a bound on the number of losses and gains of each mutation. Then we have shown how to extend it for solving the Variant Allele Frequency Factorization Problem under those evolution models.

We have performed an experimental analysis on simulated and real data which shows that the Dollo(1) (*i.e.*, the Persistent Phylogeny) and the Dollo(2) models can outperform the Perfect Phylogeny model, by measuring how close our predicted frequencies are to the measured (input) frequencies. Our ILP formulation has not been optimized for efficiency. Still, we are able to manage datasets with 20 mutations, which is common for liquid tumors. On the other hand, we need to further investigate how to extend our approach to larger instances (more samples and mutations): this will require to improve the computational efficiency of the ILP formulation or adopting some combinatorial strategies to govern the introduction of a small number of mutation losses and gains in the solution.

Finally, our comparison between our predictions and the phylogenies in the literature shows that we are abe to confirm the driver mutations or at least most of the main lineages of the trees.

## Acknowledgements

The authors would like to thank Simone Zaccaria for the useful discussion on the VAFFP and ILP formulation. We also wish to thank the reviewers for their valuable comments allowing us to improve the clarity and content of the paper.

We acknowledge the support of the MIUR PRIN 2010–2011 grant “Automi e Linguaggi Formali: Aspetti Matematici e Applicativi” code 2010LYA9RH, of the Cariplo Foundation grant 2013–0955 (Modulation of anti cancer immune response by regulatory non-coding RNAs), of the FA grants 2013-ATE-0281, 2014-ATE-0382, and 2015-ATE-0113.

